# Evidence of elemental encoding at the olfactory periphery

**DOI:** 10.1101/2025.04.21.649882

**Authors:** Daniel A Raps, Georgia M Pierce, Giulia Papiani, Lily Wu, Randy Arroyave, Jessica H Brann, Patrick Pfister

**Affiliations:** dsm-firmenich, 250 Plainsboro Road, Plainsboro, NJ 08536, USA

## Abstract

How complex, multifaceted perceptual odor qualities of volatile chemicals are encoded by the olfactory system is poorly understood. We explore the receptive fields of individual odorant receptors and show how potent agonists share a consistent perceptual quality despite chemical diversity. This supports the idea that the olfactory neural connectome maintains peripheral information like odor quality through the distinct layers of the system, akin to a labeled line regime present in other sensory structures. We further observe that simple combinatorial activation of odorant receptors can generate a perceptual sum. Human olfaction may thus be shaped by the orthogonal and elemental encoding of olfactory features detected by receptor selectivity and activity. These results provide novel insights into how the brain processes olfactory information and more broadly into neural encoding during sensory processing.

## Introduction

The olfactory system enables discrimination and recognition of an exceptionally variable and complex odorous environment. This dual function allows for odor categorization as a measure of sensitivity, intensity and quality to effectively reduce the high-dimensional odorant volatile space into a lower-dimensional and generalizable perceptual space, and provide environmental and behavioral relevance. These olfactory attributes must first be encoded in the periphery where combinatorial activity of olfactory sensory neuron (OSN) ensembles represents the gateway to perception. In mammals, each mature, synaptically integrated OSN expresses only one of a large repertoire of odorant receptors (ORs) ^1–3^, ensuring a unique molecular identity for a given neuron ^4–7^. Volatile compounds in turn exhibit a wide range of affinities for a subset of ORs and thus activate ORs selectively. The resulting receptor promiscuity leads to unique and discrete neuronal activity patterns for virtually each compound or mixture thereof ^8–11^. Our sense of smell emerges from these combinatorial representations. While distinct activity patterns may underlie the discrimination power of the olfactory system, how the brain assigns qualitative and hedonic values to odors is not fully understood. Attempts at linking chemical structure to odor percept led to mixed results in part because chemical similarity is not a rigorous predictor of perceptual similarity ^12–18^. Most complex or natural odors are comprised of a set of odorants, some of which are monomolecular chemicals. Each monomolecular chemical is not typically described by a single olfactory descriptor but rather by a set of descriptors, such as *earthy*, *musty*, and/or *green*. Such multifaceted tonalities exhibited by a single odorant have confounded the interpretation of the encoding role of each receptor bound and activated by such an odorant. Also, and often overlooked, real world olfactory percepts arise from complex mixtures that generate widespread peripheral modulation, essentially disrupting a singular link between chemistry and perception ^19–22^. In addition, olfactory bulb (OB) interneuron inhibition and cortical feedback further modify the peripheral input compared to the bulbar output ^23,24^, pointing to the importance of considering the global structure of the olfactory system to comprehend quality encoding. Its highly conserved anatomy across vertebrates and non-vertebrates suggests that the multilayered architecture is of functional relevance ^25–27^.

The molecular logic of odor encoding in the peripheral olfactory system is preserved as axonal projection emanating from OSNs expressing the same OR converge into functional units called glomeruli in the OB ^28–31^. Within each OR-specific glomerulus, axons synapse with mitral and tufted (M/T) second-order neurons. The dendritic ends of M/T cells contact only one glomerulus, thus maintaining segregation of the peripheral input ^23^. M/T cells contacting the same glomerulus also exhibit synchronized neuronal output at higher stimulus concentrations (i.e. strong agonism) thus maintaining functional encoding lines ^26,32–34^. The multiple mitral cells contacting the same glomerulus project to distributed ensembles of glutamatergic neurons of the piriform cortex (PC) that vary upon repeat odor presentation ^35^. Theoretical work shows however that this apparent representational drift is still capable of exhibiting a degree of output consistency capable of preserving qualitative peripheral information in evolving contexts ^36,37^. Overall, such functionally discrete and stereotypical neuronal wiring maintained through the distinct olfactory layers presents the outline of a labeled line signal integration. Although this structural regime does not exclude the tenets of pattern theory, we provide evidence for such labeled coding lines in the olfactory system with a quantifiable link between odorant receptor activation and semantic perceptual descriptors. Odor percepts were however reported to arise from configural neuronal representations in cortical areas such as the PC ^14,38–40^, leaving limited room to an elemental coding logic. Despite the consensus around configural encoding, evidence for elemental olfactory perception has been reported ^41^. Here we challenge the notion further that olfaction is solely configural by reporting how a subset of human ORs singularly encode individual olfactory notes, prerequisite of an elemental coding logic of odor quality.

## Results

### Human OR11A1 agonists exhibit chemical diversity and an underlying perceptual unity

Odorant receptors bind to a multitude of distinct molecules to be able to detect the sheer diversity of odorant chemicals surrounding us, an apparent promiscuity that is balanced by a high degree of receptor binding selectivity essentially segregating the detection of volatile compounds ^42^. Functional isomers or enantiomers are among the clearest example of chemical similarity ORs readily discriminate ^22,43,44^. To explore the role of a single OR within the global combinatorial olfactory activity framework, we set out to measure both physicochemical and perceptual diversity represented by the receptive field of individual receptors, including select isomers when possible. We performed cell-based dose-response assays of human ORs with volatile compounds for which sensory descriptions were available (see Methods). As an example, human OR11A1 typically exhibits both breadth and selectivity of chemical diversity. Terpene derivatives ((2)-Ethyl fenchol), sesquiterpenes (patchoulol), tertiary alcohols ((2)-Ethyl fenchol, patchoulol and (+/-)-Geosmin), or polycyclic aldehydes (trans-vulcanolide) present diverse chemistry and structures able to generate strong activation of the receptor over a range of affinities yet belong to distinct olfactive families: MUSK (trans-vulcanolide), WOODY (patchoulol), EARTHY ((+/-) geosmin). Conversely, the receptor readily discriminates the structurally and perceptually related compounds trans-vulcanolide and tonalide (full vs. partial agonist, respectively), both of which belong to the musk family. Beyond structural diversity, compounds that activate the receptor strongly share one of their olfactive facets: the earthy quality (Figure 1A, B). Structure, composition (monomolecular or mixture), and olfactive description of all agonists are listed in Supplemental Table 1. Comparing dose-response curves of structural analogs’ and isomers’ activity reveal structural constraints underlying OR11A1’s selectivity and how the stronger agonists exhibit an earthy descriptor regardless of chemical distance (Figure 1C, D). Overall, agonists exhibited a larger diversity in perceptual space (mean similarity = 0.22, Figure 1E) compared to chemical space (mean similarity = 0.46, Figure 1F). When considering the top 11 agonists only however, perceptual similarity increases dramatically (mean similarity = 0.45, Figure 1E, red box), compared to chemical similarity (mean similarity = 0.53 Figure 1F, red box), in agreement with the observation that top activators share a common descriptor.

**Figure 1:**
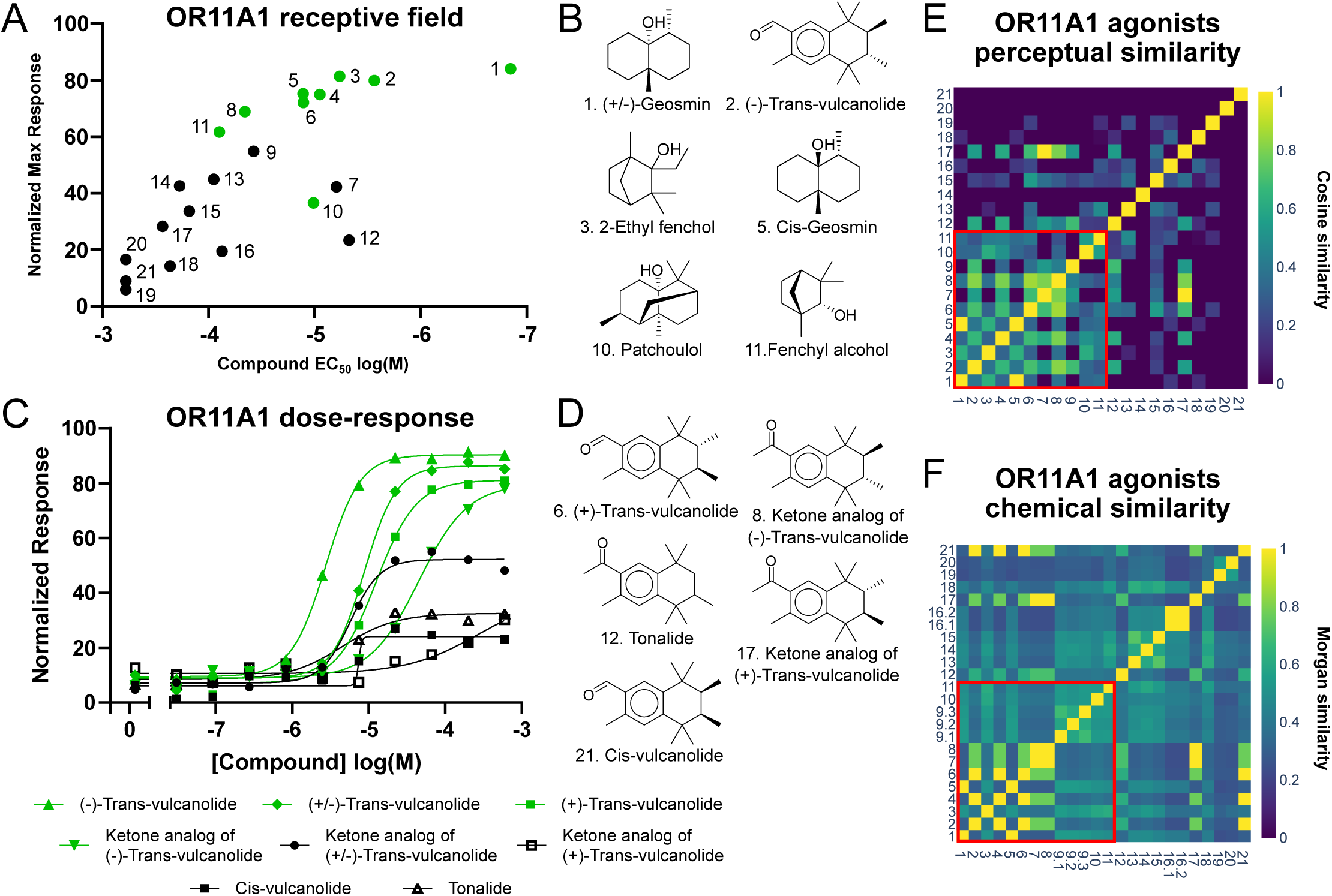
Correlation between the activation of OR11A1 and the presence of the earthy quality. (A) response OR11A1 activation profile with a panel of earthy (green dot) and not earthy (black dot) compounds (see Supp. Table 1 for complete data set, compound numbering kept consistent throughout the figure). Y axis: efficacy by homogeneous time resolved fluorescence (HTRF) maximal dose-response values, normalized to the maximal HTRF ratio induced by forskolin. X axis: potency measured by EC_50_ in log Molar. (B) Structural diversity among OR11A1 top agonists. (C) Response of OR11A1 to increasing doses of vulcanolide, tonalide and structurally related isomers and analogs. Green curves represent responses of earthy compounds. Y axis: efficacy by HTRF values normalized to maximal forskolin response (not shown). (D) Structural similarity between full and partial OR11A1 agonists. (E) Perceptual cosine similarity matrix. Similarity: mean = 0.22, top 11 agonist = 0.45 (red box). (F) Chemical Morgan similarity matrix. Similarity: mean = 0.46, top 11 agonists = 0.53 (red box).

### Evaluation of perceptual unity underlying odorant receptor activity

To evaluate the receptive field of distinct ORs as a function of perceptual diversity, we calculated the Bayesian probability of a descriptor being associated with a volatile compound as a function of receptor activation level. We standardized the activity measurements to capture both the potency and the efficacy of a receptor-ligand pair using an adapted activity index (see Methods, Figure S1), where inactive ingredients have a value of 0. A series of activity index threshold values were then selected, and at each value all ingredients for a given receptor with an activity index above threshold were classified as “activators”. This binary classification at each level allowed us to model the probability of any given olfactory descriptor being present above the threshold value as a beta distribution. An activity index threshold of 0 was included to indicate the set of ingredients that were screened in dose response against the receptor (see Figure S2). This is an important aspect of the analysis, as it demonstrates the bias of the dataset towards certain descriptors. The set that was assayed against each receptor was actually a subset of a larger initial probe of the receptors, taking advantage of the unique chemical library available to us, which covers most major perfumery families. Any chemicals from this larger set of over one thousand ingredients that passed the initial screen were moved forward for the dose response set, further biasing the input data towards molecules likely to have some level of interaction with the receptors. Despite all of this, we observed that the screening set typically showed relatively low probabilities for containing any specific descriptor. When looking at the probability of the representative descriptors EARTHY, WOODY, and MUSK associated with OR11A1 agonists, the probability of EARTHY increases as the activation threshold becomes more stringent, and converges towards certainty beyond the activity index of 1.0 (Figure 2A). For WOODY the opposite relationship is observed and compounds with higher activity index values appear to be inversely correlated to OR11A1 activation (Figure 2B). No significant shifts were observed for MUSK when comparing to the screening set (activity index threshold of 0) (Figure 2C). The same approach on the receptor OR7A17 revealed a convergence onto the WOODY tonality, a slight anti-correlation with EARTHY, and only a slight increase in probability for MUSK (Figure 2D-E). We then looked at all descriptors that appear with any ingredient screened against the receptors and plot the probability as the mean of the beta distribution, across all the threshold values. For both OR11A1 and OR7A17, we observed a strong probability of a single tonality (Figure 2G-H). We create a system-wide view with a subset of 8 receptors’ activation data and associated tonalities, by finding the top 3 descriptors and probabilities for each receptor at each previously selected activity index threshold. For each descriptor-receptor pair from this process, we find the highest probability seen, the activity index it was observed, and at how many cutoff values that descriptor was a top 3 term. We then created a matrix showing how these tonalities correlated with these receptors (Figure 2I, Table S2). A single tonality emerges for each OR: OR11A1/earthy, OR1N2/musk, OR5AN1/musk, OR5AU1/coconut, OR7A17/woody, OR8B3/tonka (a hay-like scent best represented by coumarin), and OR8D1/maple, except for OR5A1 associated with both violet and woody (see Figure 4 however). The apparent functional redundancy between OR1N2 and OR5AN1 may point to a semantic limitation of this study in capturing the granularity behind the term musk and the different types of musk ingredients. Interestingly however, there was no agonist overlap between the two receptors suggesting that more refined descriptors may help in distinguishing the receptors’ organoleptic properties. Additionally, we also observed ingredients described as earthy, musky or woody that were not found to activate the ORs tested here, indicating that these may be sufficient to elicit the corresponding olfactory responses but not required.

**Figure 2:**
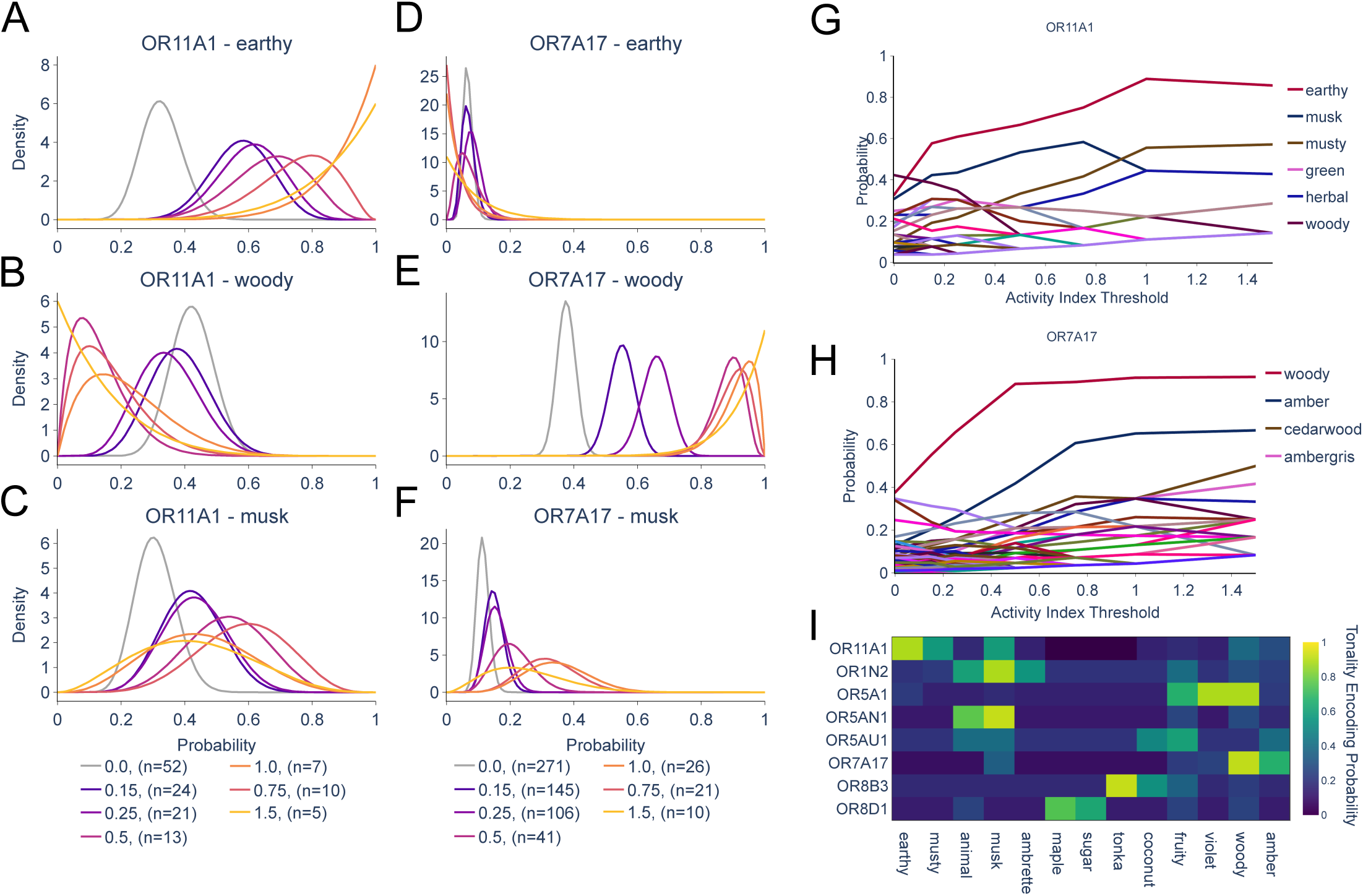
Olfactory receptor activity correlates with perception of a narrow range of tonalities. (A-C) Probability density functions of 3 separate descriptors at different activity index thresholds for OR11A1. (D-F) Probability density functions of 3 separate descriptors at different activity index thresholds for OR7A17. (G-H) Mean values of beta distributions for all present descriptors at increasing activity index thresholds for OR11A1 (G) and OR7A17 (H). (I) Heatmap of receptor-tonality probabilities, derived from the highest mean probability at any threshold value.

To further probe these receptor/tonality associations, we explored if we could predict relevant levels of activation of a receptor by providing solely the weighted descriptors for a compound. We labeled compounds as activators if the activity index they elicited was equivalent to 20% or more of the maximum activity index observed for that receptor. While many ingredients do not activate receptors by a significant amount and fall below this threshold (Figure S2), this criterion is relatively permissive and does not limit ingredients to only the highest levels of receptor activation (Figure S1). For each OR, we trained a random forest classifier to predict whether compounds activated the receptor or not based on the compound’s perceptual descriptors and their attributed weight. The models were able to accurately predict receptor activation when given the descriptors and weights (average F1 scoreof 0.98, as assessed by repeated stratified K-fold cross validation). Next, so that we might take advantage of all our data, we retrained the models on the entire dataset for each receptor. From these full data models, we asked which descriptors each model used to determine if a compound is an activator. For each receptor model, we took an explainable machine learning approach and calculated the SHAP values ^45^ for each compound. We took the average absolute value across compounds to determine the feature importance of each descriptor. This analysis revealed comparable top descriptors for each receptor as the probabilistic method (Figures 2I, 3E, Table 2). For example, the OR11A1 model listed EARTHY, WOODY, and MUSK as the top descriptors in that order (Figure 3A,C), while the probabilistic method listed EARTHY, MUSK, and MUSTY as the top descriptors. Both methods selected WOODY as the top descriptor for OR7A17 (Figure 3B).

**Figure 3:**
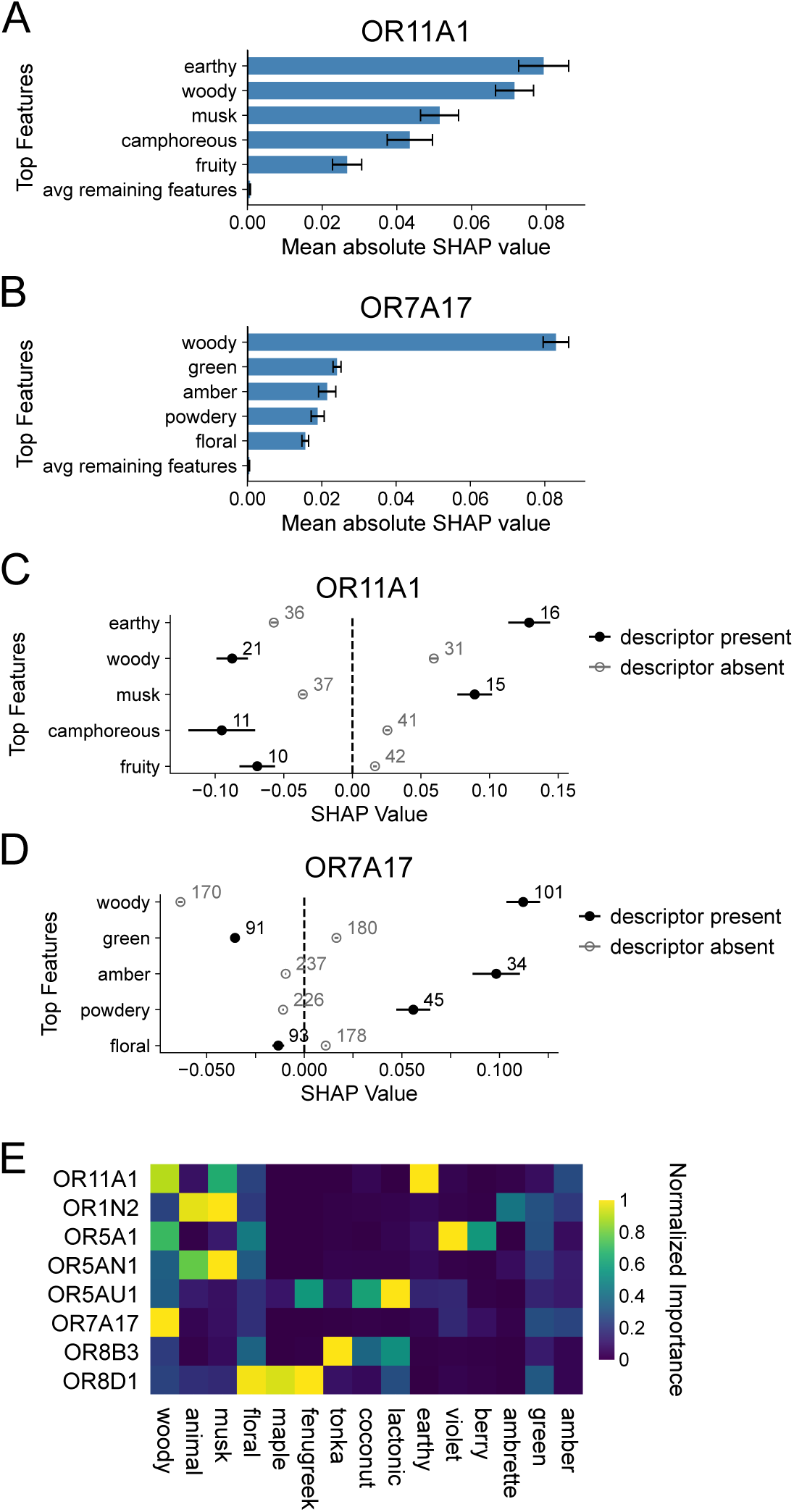
Olfactory receptor activity correlates with perception of a narrow range of tonalities. (A-B) Mean absolute value of SHAP values for the top descriptor features for the OR11A1 model and the OR7A17 model. (C-D) For the OR11A1 and OR7A17 models, average SHAP values for compounds that are described by a given descriptor (black) and those that are not (gray), error bars indicate standard error of the mean, numbers indicate the number of compounds included in each datapoint. (E) Normalized feature importance as calculated by mean absolute value of SHAP values for the top three descriptor features from each receptor model, scaled between 0 and 1 for each receptor.

**Figure 4:**
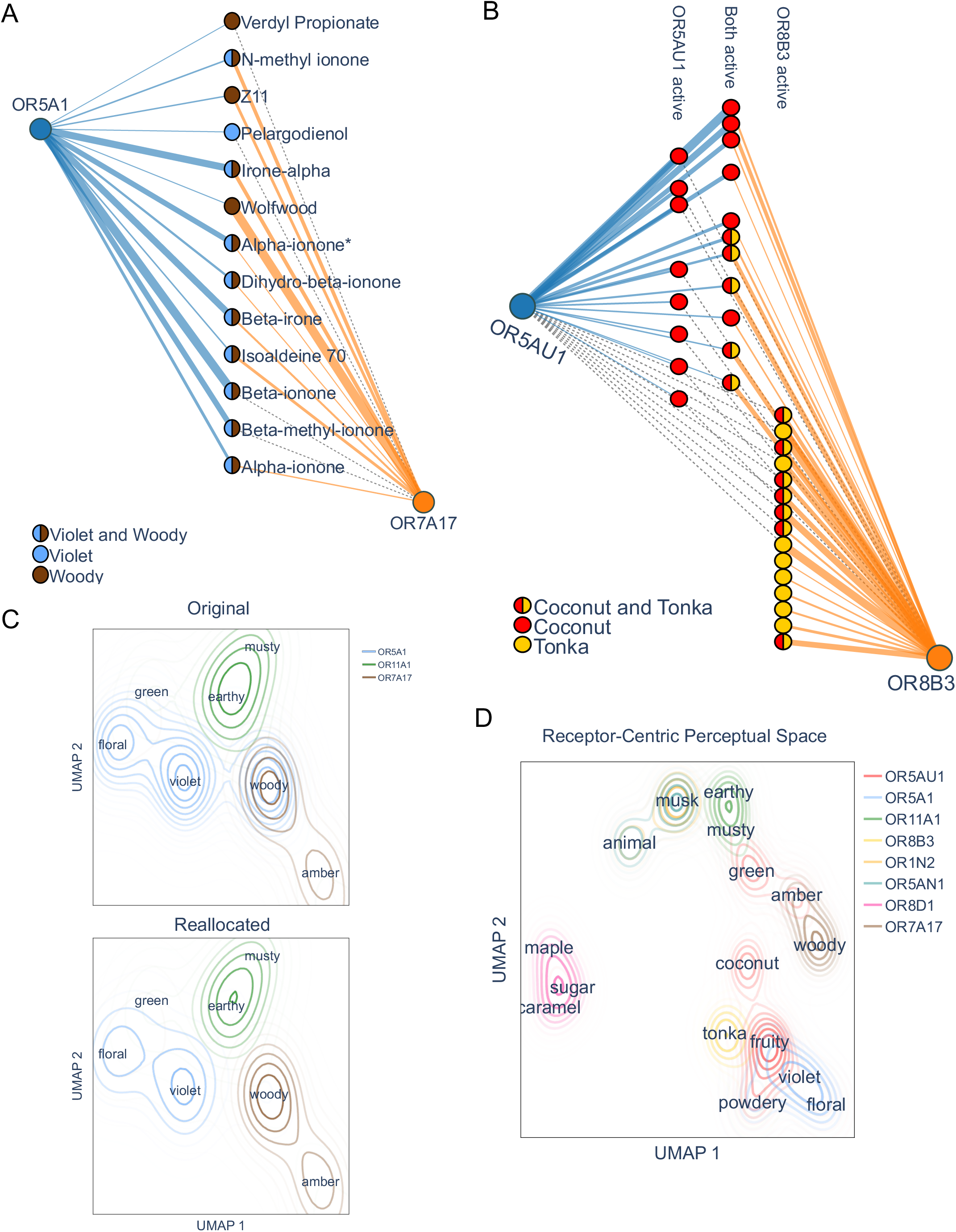
Ingredients activating multiple receptors clarify receptive range and suggest elemental encoding. (A) Network plots of ingredients activating multiple receptors. OR5A1 and OR7A17 agonists described as WOODY or VIOLET, and their corresponding activity are shown. *Denotes mixture of stereoisomers. (B) Activity network plot for OR5AU1 and OR8B3 agonists described as COCONUT/LACTONIC or TONKA/HAY. Edge thickness represents activity index between ingredient and receptor, with grey dotted lines indicating a pair was screened but showed no activity. (C) Dimensionality reduction of perceptual space, built from descriptor cooccurrence in ingredients with contour lines of receptor probabilities, according to kernel density estimates, showing original probabilities (top) and after running the reallocation algorithm (bottom). (D) Receptor-centric perceptual space, constructed from the reallocated probabilities matrix of receptor and tonalities.

One advantage of analysis with SHAP is that SHAP values are designed to allow for inspecting the model decisions of individual inputs/compounds in addition to the feature importance of descriptors as a whole. We decided to investigate how ingredients might differentially impact model decisions about receptor activation if they are described as having a certain olfactive descriptor or not. We took the average SHAP value for compounds that had positive weights for each descriptor and the average for ingredients that were not described as having each descriptor. We saw that for compounds described by the key tonalities for each receptor, such as EARTHY for OR11A1 or WOODY for OR7A17, the average SHAP values were in the positive range. This result indicates that the model was more likely to classify these compounds as activators of the receptor (Figure 3C,D). Conversely, we saw that compounds that were not described with the key descriptor (EARTHY for OR11A1 or WOODY for OR7A17) had negative averages. The model was less likely to designate those compounds as activators. Thus, the OR11A1 model learned that earthy-smelling compounds are more likely to activate OR11A1 and compounds that do not smell earthy are less likely to elicit much activation from OR11A1, in agreement with the results in Figure 1. This analysis also corroborates the anti-correlations discovered by the probabilistic method. Compounds described as woody tended to have a negative SHAP value in the OR11A1 model, demonstrating a lower likelihood of being classified as OR11A1 activation. Compounds not described as woody were more likely to be classified as activators of OR11A1. Similarly, compounds described as green tended to have a negative SHAP value in the OR7A17 model, suggesting a lower likelihood of being labeled OR7A17 activators. Conversely, compounds not described as green had a SHAP value in the positive range, indicating that they are more likely to be classified as activators of OR7A17, though the value is small. These explainable machine learning techniques support the link between OR activation and olfactory tonality perception. By examining SHAP-based feature importance of the random forest models of each receptor, we were able to corroborate the findings from the probabilistic analysis and assign linked tonalities to each receptor. These links become even clearer when examining the average SHAP values for compounds positive for those tonalities, such as where we observe woody and amber ingredients demonstrating higher likelihood of being classified as OR7A17 activators (Figure 3D).

### Co-activation of odorant receptors reveal odor encoding elemental in nature

In Figures 2 and 3, we evaluated how OR activities are correlated with distinct perceptual features and showed that OR7A17 is strongly correlated with WOODY, and how OR5A1 exhibits high probabilities for both VIOLET and WOODY. When we examined the individual ingredients that are activating OR5A1 however, we saw that many of them are in fact co-activating OR7A17 (Figure 4A). For each receptor, we gathered a list of the most likely descriptors from the original probabilistic association approach. We then recalculated the probabilities for each receptor-tonality pair, with the condition that if an ingredient contains a descriptor that is linked to another receptor, and also activates the other receptor, that descriptor gets allocated to its linked receptor and removed from other receptors. If the descriptor is linked to another receptor, but the ingredient does not activate the linked receptor, then it is included in the calculations for the receptors it does activate. When we allocate the “woodiness” of those ingredients to OR7A17 activation instead, the probability of OR5A1 being linked to WOODY falls and we are left with a clear link to VIOLET and the suggestion of an additive encoding strategy.

We then performed this type of reallocation for all of our tonality-receptor relationships. Even with this reallocation, the receptors OR5AU1 and OR8B3 appeared to be quite overlapping in their tonalities. To investigate this, we selected all ingredients that activate either of these receptors at an activity index of 0.25 or greater, and contain any of the descriptors COCONUT, LACTONIC, TONKA, or HAY. We then viewed the ingredients and receptors as a connected network and observed that the OR5AU1 agonists lean COCONUT, whereas TONKA aligned with OR8B3 activation (Figure 4B). Examining these pairs of terms highlighted an obstruction to this type of work, namely the gulf between semantic availability and sensory experience. Often, HAY and TONKA may be used interchangeably when discussing scents, however that leads to lower overall probability of each term individually, so these terms were merged for this approach. Likewise, COCONUT and LACTONIC notes share a certain aspect in olfaction, making disentanglement between the terms an impediment to this type of work. It is possible that OR5AU1 is actually related to the COCONUT side of lactones, or the LACTONIC side of coconuts, and not the merging of the two terms as we have performed here. When we viewed the ingredients and their descriptors from the perspective of their receptor activity, we saw what appeared to be perceptual summation, rather than synergy. For example, activating OR5AU1 alone was likely to elicit just COCONUT, and OR8B3 activation alone lead to a TONKA percept. The ingredients that activate both, however, are likely to be perceived as both coconut and tonka.

We constructed a perceptual space by running a dimensionality reduction algorithm on the co-occurrence of descriptors across the entirety of the data set, and then using the Bayesian probabilities as an input for kernel density estimates to create a contour map of the receptor coverage of olfactory perceptual space. Examining the region of space surrounding the terms important to OR5A1 and OR7A17 demonstrates the effect of our reallocation method (Figure 3C). In the original space, OR5A1 covers a broad range of both VIOLET and WOODY, however after the reallocation process, OR5A1 is strongly centered on VIOLET and OR7A17 corresponds to WOODY. For both receptors we saw secondary clusters, OR5A1 around FLORAL, possibly due to an issue in descriptor granularity, and OR7A17 around AMBER which may suggest co-activation with additional receptors that were not examined here. Finally, we created a receptor-centric view of olfactory perceptual space by performing the dimensionality reduction on the reallocated matrix of probabilities. The resulting space showed distinct regions of percepts as they are viewed through the lens of receptor activity (Figure 4D).

## Discussion

Here we report how a single common olfactive descriptor is observed amongst the strong agonists of individual ORs independently of the chemical structure and the overall multifaceted descriptive nature of odorants, including monomolecular volatiles. Within receptive fields, major agonists share an underlying tonality regardless of chemical diversity, a phenomenon that is not observed when randomly sampling volatiles from the large set representative of odor space considered here (Figure S3). Similarity in organoleptic properties (i.e. receptor affinities linked to perceptual outcome) thus translate into a perceptual common denominator for compounds that share their affinity to specific ORs. Human perceptual representation of volatile chemical environment may thus be confined by the orthogonal encoding of olfactory features detected by receptor selectivity and activity, and by extension OSNs that express them, Interestingly, a similar activity segregation pattern was reported across the olfactory repertoire of the fly larva. Increasing concentration of distinct volatile compounds led to the orthogonal recruitment of olfactory receptor neurons. These compounds all have distinct smells and are likely to represent different environmental cues ^46^.

The apparent discrepancy between chemical and perceptual similarity can now also be considered under the lens of OR activity as a better predictor of olfactory percept compared to chemical structure alone^47^, providing receptor-driven structural constraints for the discovery of new perfumery ingredients. This implies that the olfactory neural connectome can maintain peripheral information such as olfactive quality segregated through the distinct layers of the system, akin to labeled lines seen in other sensory systems. This is supported by studies observing how intensity and quality may be encoded by distinct cortical neuronal population as suggested by work in mice and humans ^41,48,49^. The potential role of a labeled line for more downstream odor-related percepts, such as valence or contextual interpretation, remains however questionable as these involve more complex central integration circuits ^50^. As opposed to the emergent sensory properties that can emanate from complex codes ^51^, elemental encoding calls for additive effects where the combination of multiple receptors generate a perceptual sum rather than a synergy. Despite a relatively small ligand-receptor pair sample size, we in fact observed several cases of compounds described by multiple tonalities activating ORs whose attributed tonalities corresponded. Such a coding strategy may thus play a role in the perceptual processing of odors, notwithstanding the inherent subjectivity of the olfactory sensory modality influenced by cultural, semantic ^52,53^, and genomic diversity encountered in humans

Such a coding strategy may be limiting for highly complex odor environment but sufficient for simpler conditions that derive from the olfactory system’s tendency to maintain encoding sparsity. In rodents for example, individual odorants have been shown to elicit sparse activation codes in the OB when facing low but realistic concentration ranges, activating their most sensitive respective ORs ^54^. In addition, at each layer of the olfactory system, specific processes improve sparsity further. In the periphery, code saturation is limited by widespread inhibition when presented with odor mixtures ^19–21^. In the bulb, gain control and figure-ground separation occur by lateral inhibition as well as cortical top-down negative feedback from the piriform cortex ^24^. An elemental encoding logic may thus play a predominant role as a function of high sparsity to encode multifaceted olfactory percepts.

## STAR+METHODS

Detailed methods are provided in the online version of this paper and include the following:

### KEY RESOURCES TABLE

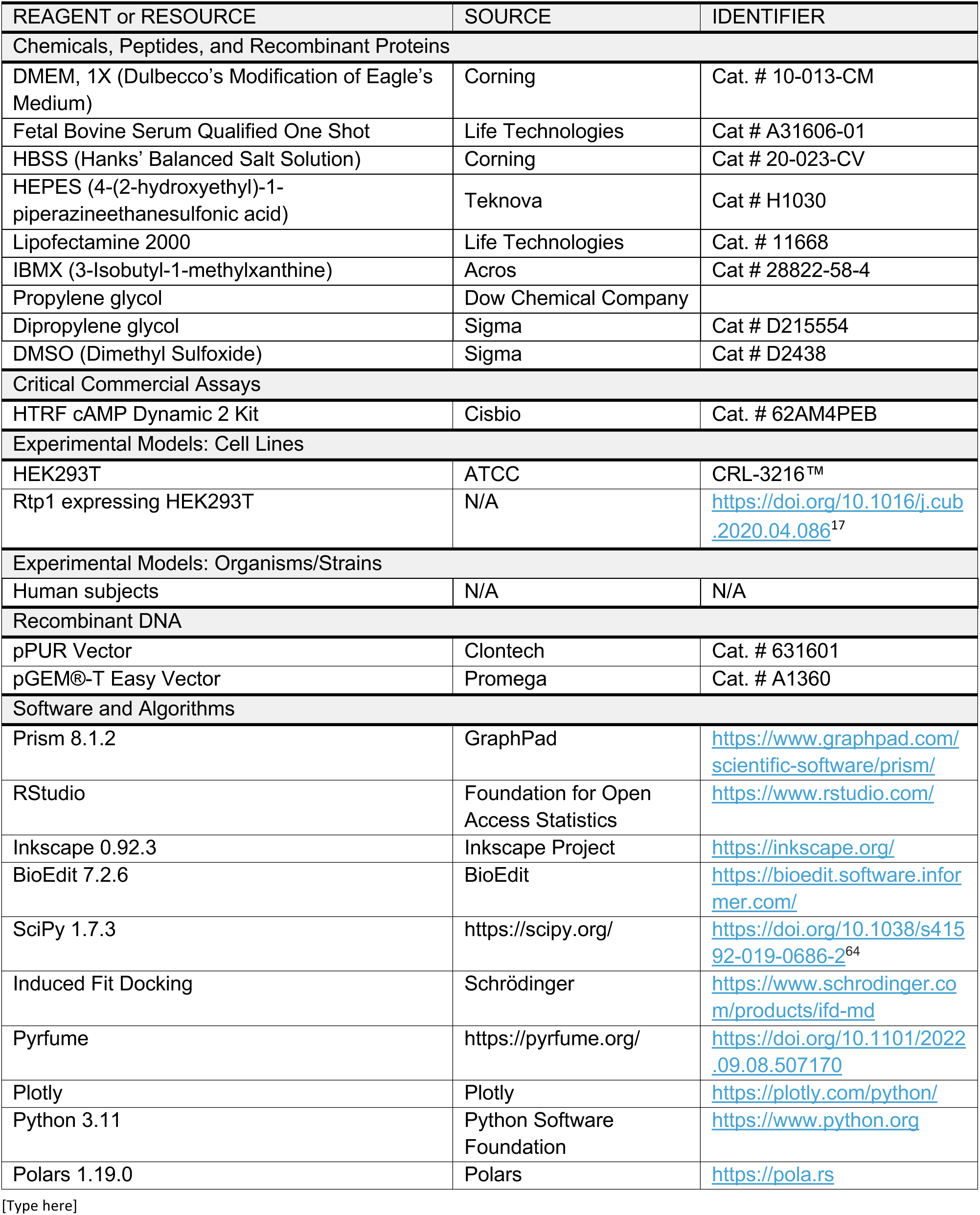
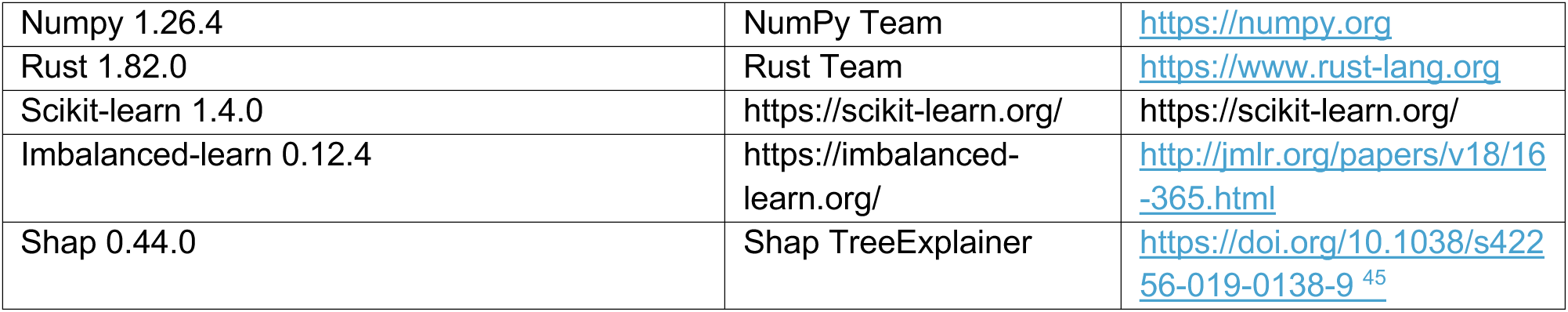

| REAGENT or RESOURCE | SOURCE | IDENTIFIER |
| --- | --- | --- |
| Chemicals, Peptides, and Recombinant Proteins |  |  |
| DMEM, 1X (Dulbecco's Modification of Eagle's Medium) | Corning | Cat. # 10-013-CM |
| Fetal Bovine Serum Qualified One Shot | Life Technologies | Cat # A31606-01 |
| HBSS (Hanks' Balanced Salt Solution) | Corning | Cat # 20-023-CV |
| HEPES (4-(2-hydroxyethyl)-1-piperazineethanesulfonic acid) | Teknova | Cat # H1030 |
| Lipofectamine 2000 | Life Technologies | Cat. # 11668 |
| IBMX (3-Isobutyl-1-methylxanthine) | Acros | Cat # 28822-58-4 |
| Propylene glycol | Dow Chemical Company |  |
| Dipropylene glycol | Sigma | Cat # D215554 |
| DMSO (Dimethyl Sulfoxide) | Sigma | Cat # D2438 |
| Critical Commercial Assays |  |  |
| HTRF cAMP Dynamic 2 Kit | Cisbio | Cat. # 62AM4PEB |
| Experimental Models: Cell Lines |  |  |
| HEK293T | ATCC | CRL-3216™ |
| Rtp1 expressing HEK293T | N/A | https://doi.org/10.1016/j.cub.2020.04.086<sup>17</sup> |
| Experimental Models: Organisms/Strains |  |  |
| Human subjects | N/A | N/A |
| Recombinant DNA |  |  |
| pPUR Vector | Clontech | Cat. # 631601 |
| pGEM®-T Easy Vector | Promega | Cat. # A1360 |
| Software and Algorithms |  |  |
| Prism 8.1.2 | GraphPad | https://www.graphpad.com/scientific-software/prism/ |
| RStudio | Foundation for Open Access Statistics | https://www.rstudio.com/ |
| Inkscape 0.92.3 | Inkscape Project | https://inkscape.org/ |
| BioEdit 7.2.6 | BioEdit | https://bioedit.software.informer.com/ |
| SciPy 1.7.3 | https://scipy.org/ | https://doi.org/10.1038/s41592-019-0686-2<sup>64</sup> |
| Induced Fit Docking | Schrödinger | https://www.schrodinger.com/products/ifd-md |
| Pyrfume | https://pyrfume.org/ | https://doi.org/10.1101/2022.09.08.507170 |
| Plotly | Plotly | https://plotly.com/python/ |
| Python 3.11 | Python Software Foundation | https://www.python.org |
| Polars 1.19.0 | Polars | https://pola.rs |
| [Type here] |  |  |
| Numpy 1.26.4 | NumPy Team | https://numpy.org |
| Rust 1.82.0 | Rust Team | https://www.rust-lang.org |
| Scikit-learn 1.4.0 | https://scikit-learn.org/ | https://scikit-learn.org/ |
| Imbalanced-learn 0.12.4 | https://imbalanced-learn.org/ | http://jmlr.org/papers/v18/16-365.html |
| Shap 0.44.0 | Shap TreeExplainer | https://doi.org/10.1038/s42256-019-0138-9<sup>45</sup> |

### CONTACT FOR REAGENT AND RESOURCE SHARING

#### Lead Contact

Further information and requests for resources and reagents should be directed to and will be fulfilled by the lead contact, Patrick Pfister.

#### Materials availability

Volatile compounds were sourced internally, all other reagents are commercially available (see key resources table). An Rtp1-expressing HEK293T cell line was used. There are restrictions to the availability of this cell line due to existing intellectual property.

## METHOD DETAILS

### Cell maintenance and Homogeneous time resolved fluorescence (HTRF) assay

Cell-based assays were performed using modified HEK293T cells that express the Receptor-Transporting Protein 1 (RTP1) ^21^. Cells were maintained in Dulbecco’s Modified Eagle’s Medium (DMEM) containing 10% fetal bovine serum (FBS). Cells were maintained at 37°C with 5% CO2. Cells were seeded at a density of 7500cells/well in 96-well half area white plates and grown for 24 hours in media containing 10% FBS without antibiotics. After 24 hours, cells were transfected with 4.5 μg of OR and 0.5 μg of accessory protein (Golf) per plate using a standard Lipofectamine 2000 (Invitrogen) protocol. 18-24 hours post transfection, media was removed, and cells were incubated with 40 μL of compound in assay buffer (1X HBSS, 10 mM MgCl2, 20 mM HEPES, 2 mM CaCl2, 0.5 mM IBMX). Compounds were serial diluted in 100% DMSO and added to assay buffer to a 0.2% final concentration of DMSO. Cells were then incubated at 37°C in 5% CO2 for 30 minutes, lysed following the protocol for the HTRF cAMP Gs Dynamic Kit (Revvity), and fluorescence read on the PheraStar FSX (BMG LabTech). Maximum response was calculated as the percentage of the maximum window of activation for the OR, the difference between the negative (DMSO alone) and positive (forskolin) controls.

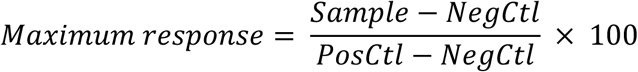

Where PosCtl = HTRF value of a forskolin dose response at saturation, Sample = HTRF value of the OR treated with odor, and NegCtl = HTRF value for DMSO alone.

### Activity index

Agonist activity levels were captured as the combination of efficacy and potency by adapting a dose response activity indexing approach ^55^. Briefly, for each compound, the maximum efficacy (E_max_) value was normalized by a factor of 100, which brings it into the same order of magnitude as the potency value. Then the negative of the difference between the log of the EC_50_ in molar and the roof of the assay (600µM, ∼3.22) was calculated, leading to an activity index of 0 for inactive ingredients.

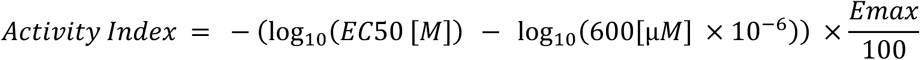

### Perceptual similarity

For a clean and unified perceptual descriptor set, the ontology publicly available in GoodScents was evaluated by internal Perfumers. Some modifications were proposed and accepted, such as merging multiple terms into a single descriptor for semantic reasons, dropping terms more related to valence, or splitting single terms into multiple descriptors for greater precision. Each descriptor was then assigned into a broad Family, as appropriate. See Supplemental Table 1 for all modifications and the final descriptor set used.

### Chemical similarity

For all of the ingredients with perceptual descriptors gathered, associated SMILES were also compiled. The chemical similarity of these ingredients were then calculated by feeding their SMILES strings into the Pyrfume package (<u>The Pyrfume Project – The Pyrfume Project – Tools, Models, and</u> <u>Data for Odorant-Linked Research</u>) to get both Morgan Features and Morgan Similarity scores.

## QUANTIFICATION AND STATISTICAL ANALYSIS

### Probability distribution functions

The probability of an ingredient that activates a receptor at or above a given activity index cutoff was modelled as a beta distribution with parameters (α, β) = 1. This sets an unbiased prior where all outcomes begin with an equal probability. The probability distribution functions were then calculated for the set of ingredients screened against the receptor (cutoff=0), and an increasingly stringent series of cutoff values (0.15, 0.25, 0.5, 0.75, 1.0, and finally 1.5).

For each receptor, at each activity index cutoff value greater than zero, the top 3 descriptors were found. This list was then filtered for descriptors that appear in the top 3 for a receptor at more than 2 cutoff levels, which lead to between 2 and 4 “top” descriptors per receptor. Each descriptor in this list was then given a probability for each receptor as the highest mean of the probability distribution seen for that tonality-receptor pair at any cutoff greater than zero. In the reallocation approach, we checked if an ingredient has the tonality assigned to another receptor, and if it activated that receptor. If so, we considered that descriptor’s presence to be due to the tonality-linked receptor, and did not count it towards other receptors’ probability calculations.

### Random Forest Models and SHAP analysis

We built and trained a random forest model to classify compounds into strong or low activators for each olfactory receptor. Input features were the weighted perceptual descriptors for the compounds tested against that receptor. The positive class consisted of compounds with an activity index equivalent to 20% or more of the maximum activity index seen for that OR. The negative class included compounds with low or no activation whose activity index was below the threshold. Given that each receptor’s data has few compounds in the positive class, activators, and many compounds in the negative class, low- or non-activators, we applied the Synthetic Minority Oversampling Technique (SMOTE) to correct for imbalanced data ^56^. This technique oversamples the positive class by generating synthetic examples that are nearby in the feature space. We employed Repeated Stratified K-Fold cross-validation to evaluate the model’s performance. The average F1 score across models was 0.98, which indicates good precision and accuracy.

After establishing the model’s prediction accuracy, we trained the models on the entire oversampled dataset. We calculated SHAP (Shapley Additive exPlanations) values for each compound using the TreeExplainer method ^45^. To determine the importance of each tonality in the model’s prediction of a given receptor’s activation, we took the average absolute value of the SHAP values across compounds for each descriptor feature. The average SHAP value was computed for compounds that were described by a descriptor and separately for compounds that were not described by a descriptor to interpret how a compound that smelled like that descriptor influenced the model to select towards or away from activating the receptor above threshold in the model’s prediction.

### Perceptual space generation

A matrix was constructed of the ingredients assayed and their perceptual description, where each row represented a single ingredient, each column a descriptor, and the cells contained the weighting of that descriptor for that ingredient. Missing values in the matrix were considered to have a value of 0. Dimensionality reduction was performed on this matrix using the uniform manifold approximation and projection method with parameters (n_neighbors=5, min_dist=0.1, metric=correlation), to create a 2-dimensional plot of perceptual olfactory space. We then limited this space specifically to the top descriptors (described above). For each receptor, we calculated a kernel density estimate on this two-dimensional space, weighted by the probabilities of each descriptor-receptor pair, and then plotted onto the space as contour lines. We also constructed a receptor-centric view of perceptual space, where the dimensionality reduction was performed on the reallocated matrix of descriptor-tonality probabilities.

## SUPPLEMENTAL INFORMATION

**Figure S1.**
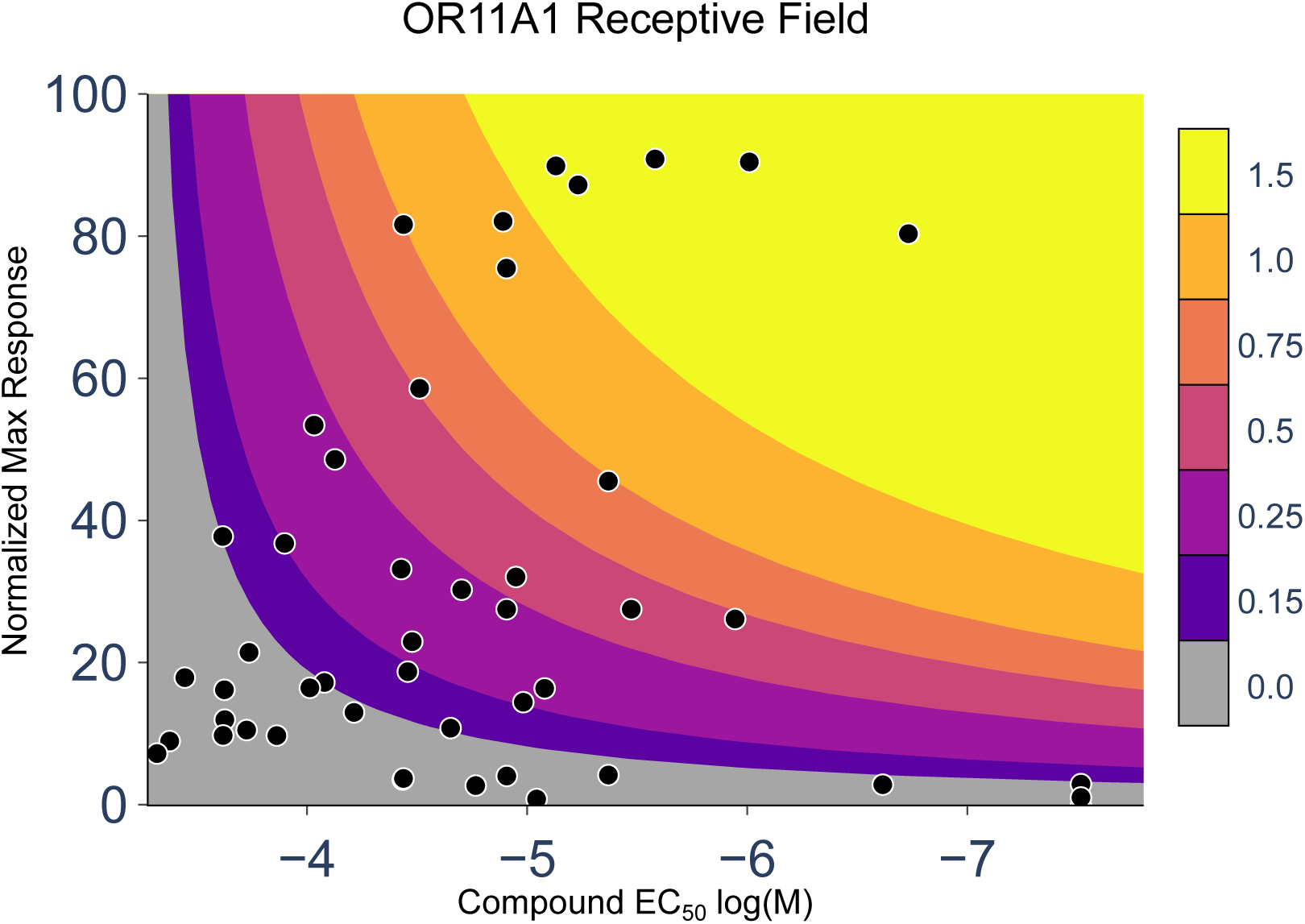
Deriving activity index from efficacy and potency: Curves show activity index equivalent of the combinations of efficacy (normalized max response) and potency (EC_50_). A single activity index can be reached with multiple different combinations of efficacy and potency. Points are values from Figure 1, with some additional points not shown in Figure 1 for clarity.

**Figure S2.**
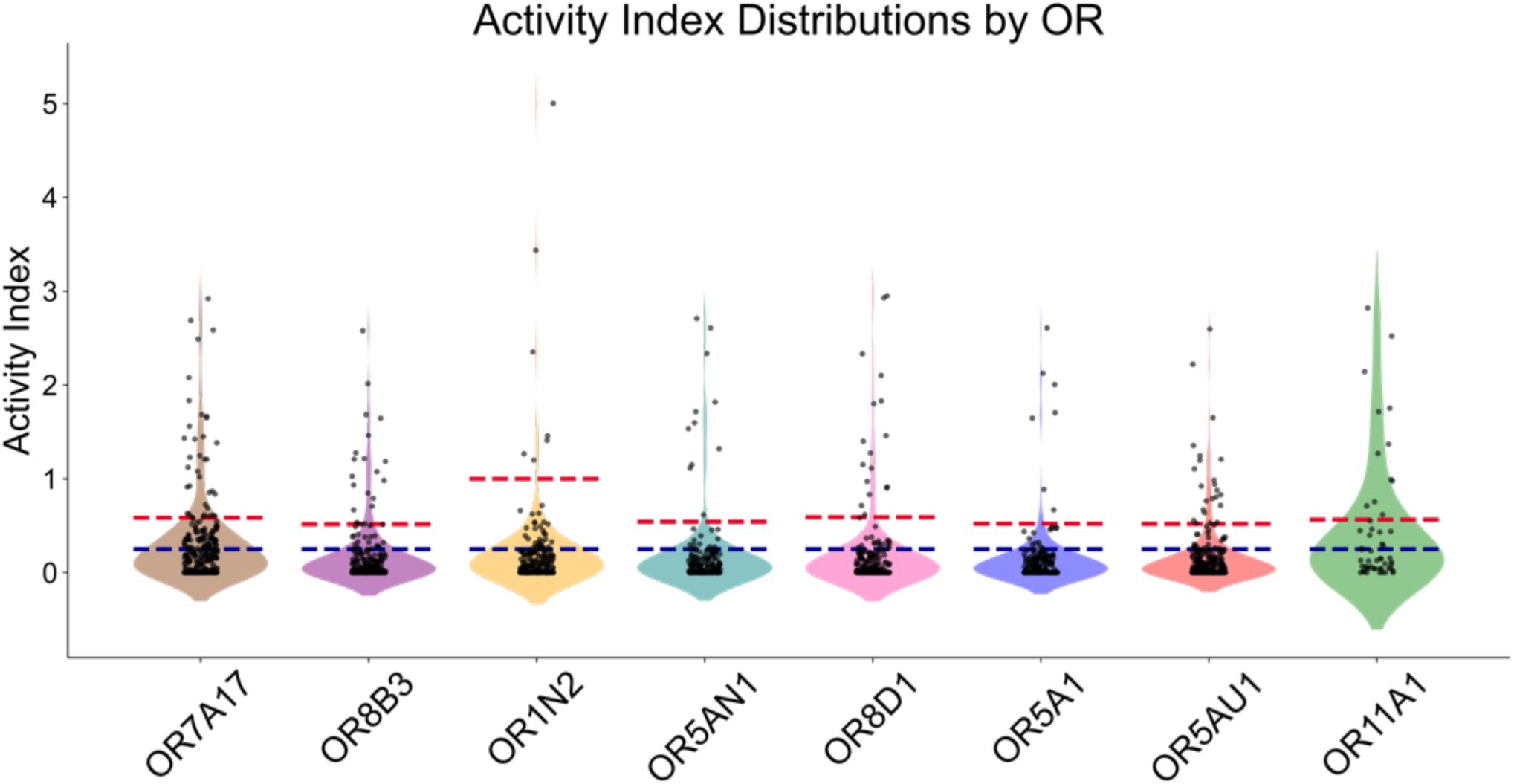
Distribution of receptor activation: Activity indices of ORs for all tested compounds. Red dotted line indicates the threshold (≥20% of max activity observed for that receptor) for classifying activators and low/non-activators in the random forest models. Blue dotted line indicates the threshold for receptor-centric perceptual space calculations in Figure 4.

**Figure S3.**
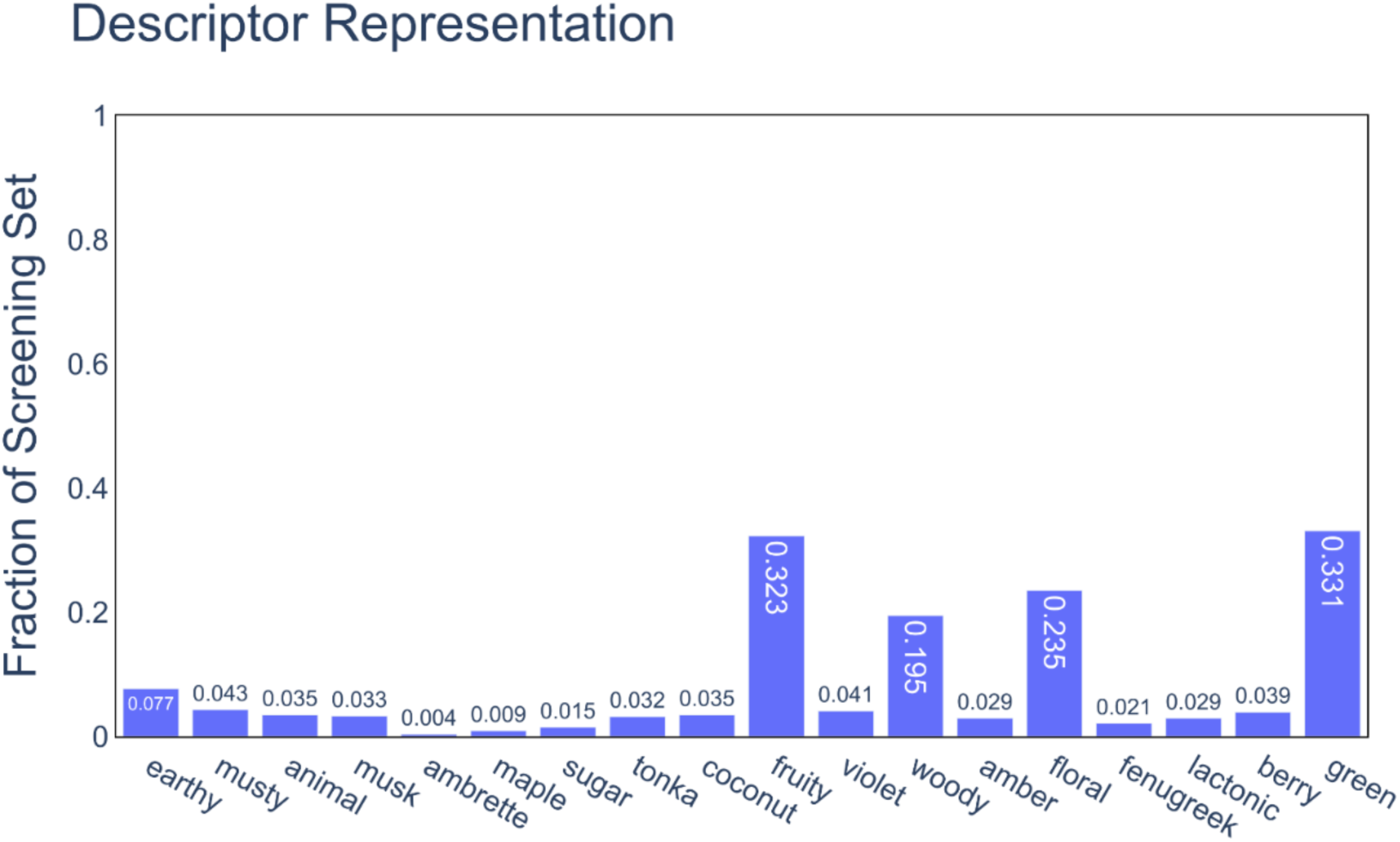
Descriptor Representation in Screening Set: The distribution of a selection of descriptors in the screening set, demonstrating the bias of the dataset. Some broad terms, such as fruity and floral, have a larger level of representation than more specific descriptors.

**Table S1.**
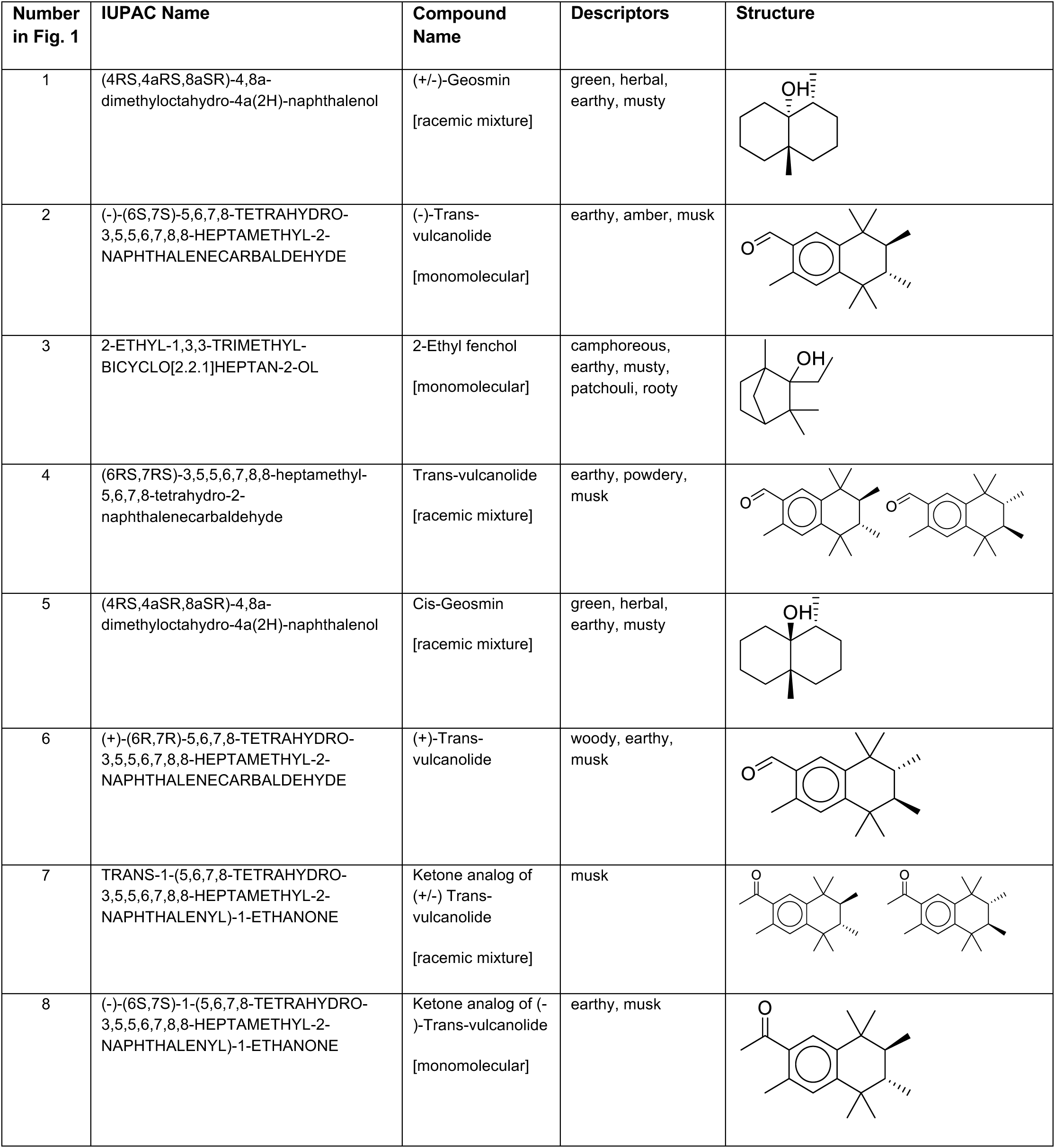
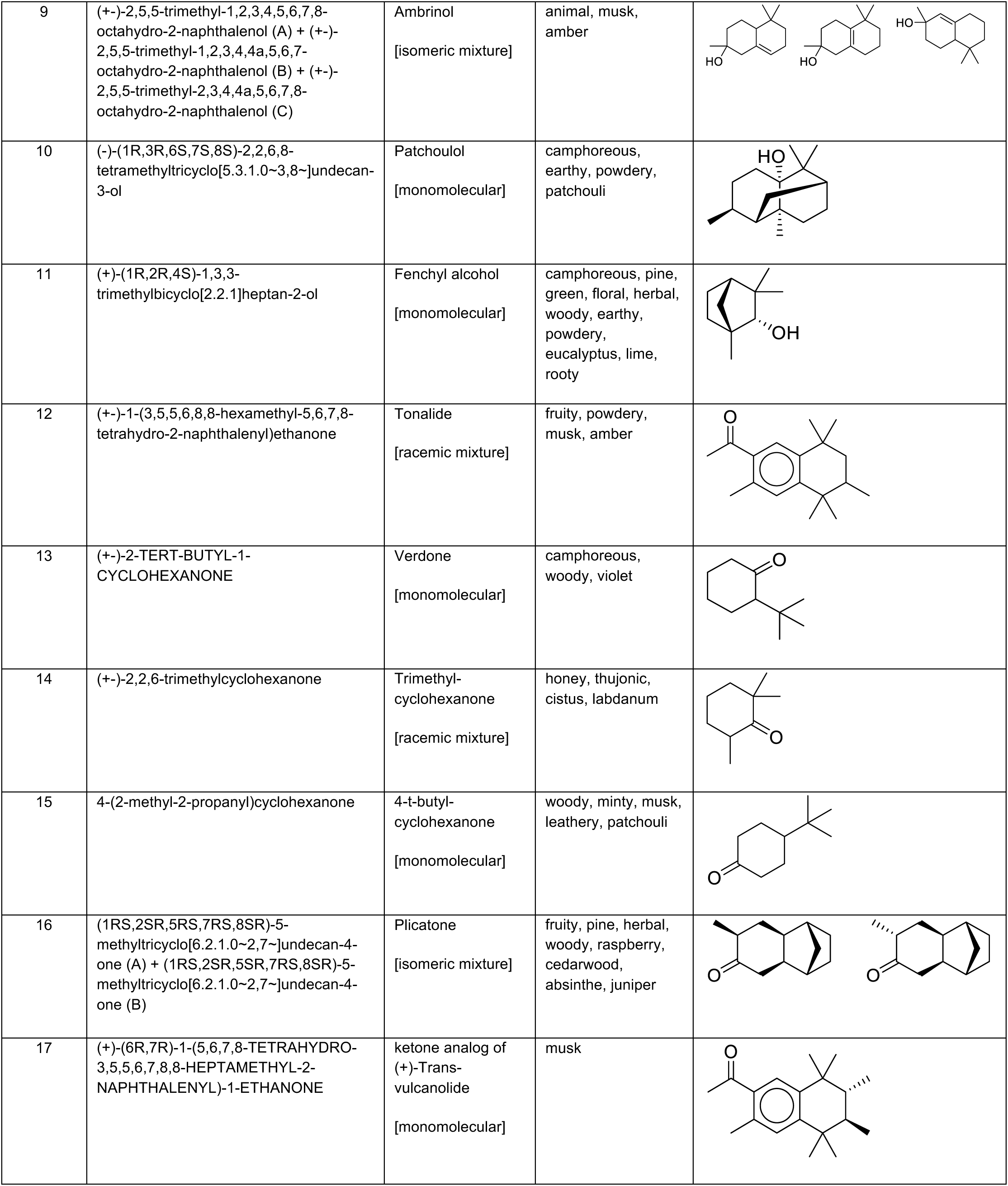
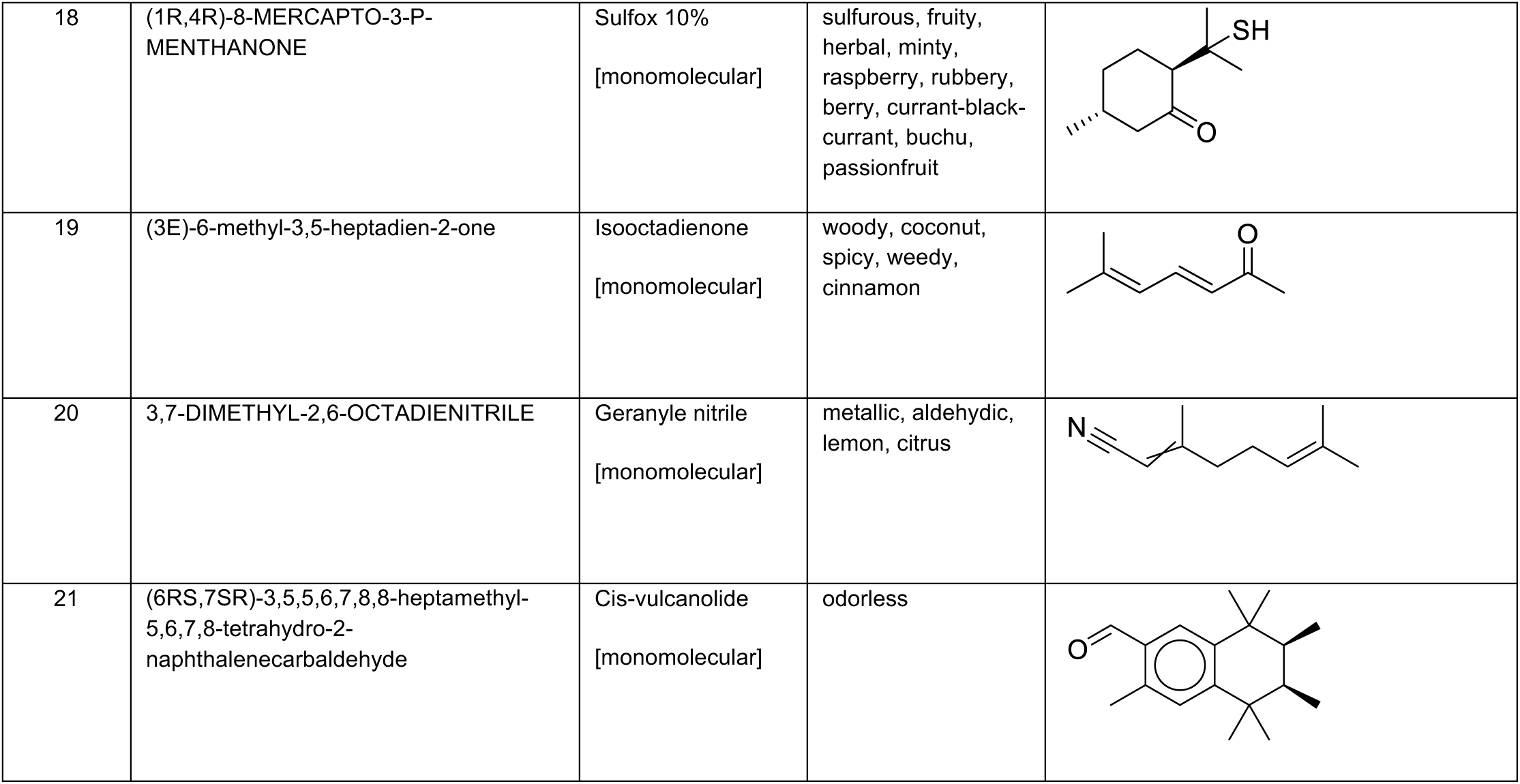
List of compounds in. Figure 1: IUPAC and common names, and perceptual descriptors and structures for the 21 compounds disclosed in Figure 1.

**Table S2.** Comparison of analyses: The top 2-3 descriptors revealed by the probabilistic method (highest mean probability at any threshold value) and the top 3 descriptors revealed by the explainable machine learning method (largest average absolute value of SHAP) for each OR.

| OR | Probabilistic Tonality | Probability | Explainable ML Tonality | Feature Importance |
| --- | --- | --- | --- | --- |
| OR11A1 | earthy | 0.8889 | earthy | 0.0793 |
|  | musk | 0.5833 | woody | 0.0715 |
|  | musty | 0.5714 | musk | 0.0514 |
| OR1N2 | musk | 0.8889 | musk | 0.0272 |
|  | animal | 0.6000 | animal | 0.0263 |
|  | powdery | 0.3571 | ambrette | 0.0120 |
| OR5A1 | violet | 0.8889 | violet | 0.0229 |
|  | woody | 0.8889 | woody | 0.0161 |
|  | floral | 0.7500 | berry | 0.0131 |
| OR5AN1 | musk | 0.9231 | musk | 0.0423 |
|  | animal | 0.7692 | animal | 0.0331 |
|  | powdery | 0.5000 | woody | 0.0147 |
| OR5AU1 | fruity | 0.6000 | lactonic | 0.0239 |
|  | coconut | 0.5000 | coconut | 0.0144 |
|  | green | 0.4444 | fenugreek | 0.0136 |
| OR7A17 | woody | 0.9167 | woody | 0.0830 |
|  | amber | 0.6667 | green | 0.0241 |
|  |  |  | amber | 0.0215 |
| OR8B3 | tonka | 0.9231 | tonka | 0.0424 |
|  | coconut | 0.5385 | lactonic | 0.0228 |
|  |  |  | coconut | 0.0161 |
| OR8D1 | maple | 0.7500 | fenugreek | 0.0181 |
|  | caramel | 0.6250 | floral | 0.0179 |
|  | sugar | 0.6520 | maple | 0.0172 |

## ACKNOWLEDGMENTS

We would like to thank members of the dsm-firmenich Science & Research division for their contributions, in particular M. Emberger for compound analysis and management and his team for the sourcing, analysis and management of the volatile compounds used in this study; Z. Peterlin and M. E. Rogers for valuable comments on the manuscript; F. De Nanteuil for chemistry insights and B. C. Smith for initial discussions.

## AUTHOR CONTRIBUTIONS

Conceptualization, D.R., J.H.B. and P.P.; methodology and investigation, D.R., G.M.P., G.P., L.W., I.A., and R.A.; formal analysis, D.R., G.M.P., G.P. and P.P.; writing – original draft, D.R. G.M.P. and P.P.; writing – review & editing, D.R., G.M.P. J.H.B., and P.P.; supervision, P.P.

